# StainX: GPU-accelerated batch stain normalization for computational pathology at scale

**DOI:** 10.64898/2026.08.06.743198

**Authors:** Samir Moustafa, Yimin Zheng, André F. Rendeiro

## Abstract

Stain normalization reduces color variability in histopathology whole-slide images, but cohort-scale pipelines lack fused multi-image batch transforms for classical methods. We present StainX, a GPU-accelerated batch stain normalization framework built around a two-stage fit/transform interface. It implements histogram matching, Macenko, and Reinhard normalizers through a portable PyTorch backend and an optional CUDA backend that fuses per-pixel operations for batch throughput. On NVIDIA GPUs, the fused CUDA path outperforms the torch CPU backend by 168×, 70×, and 48× for Reinhard, histogram matching, and Macenko respectively, and exceeds the fastest GPU peers by 7-8× (Reinhard) and 2× (Macenko) at comparable accuracy. StainX also provides user-selectable precision modes, a documented Python API, continuous integration testing, and online documentation. Source code available at https://github.com/rendeirolab/stainx, and documentation at https://stainx.readthedocs.io. Implemented in Python. Runs on Linux, macOS, and Windows.

## Introduction

Self-supervised foundation models trained on millions of hematoxylin and eosin (H&E) whole-slide images (WSIs) have transformed computational pathology, enabling general-purpose feature extraction for disease detection, subtyping, and prognostication (Chen et al. 2024). Yet despite these advances, downstream performance remains sensitive to non-biological color variation caused by differences in staining protocols, reagent batches, and scanner hardware (Tellez et al. 2019). Stain normalization corrects these artifacts by standardizing color appearance, but at the scale of modern WSIs—where a single WSI yields tens of thousands of tiles— the correction is computationally expensive. Tellez et al. (2019) measured classical stain-color normalization at 21.8–111.2 minutes per 50, 000 × 50, 000 WSI, so normalizing a large cohort can add days of compute. Normalization is therefore often skipped, leaving a known source of batch effects unaddressed in large-scale studies. When normalization is applied, it must be done carefully: a poorly chosen reference image or a mismatch in tissue composition can introduce artifacts.

Three classical approaches exist. Histogram matching aligns brightness and intensity distributions to a reference (Gonzalez and Woods 2009), Macenko normalization separates and rescales the H&E stain components (Macenko et al. 2009), and Reinhard color transfer matches overall color appearance to a reference (Reinhard et al. 2001). These methods are training-free and reference-controlled, making them well-suited for shared benchmarks, repeatable inference, and auditable cross-cohort preprocessing. Tellez et al. (2019) found that normalization alone was insufficient for top classification performance and recommended pairing it with stain augmentation. While learning-based alternatives such as StainGAN and StainNet exist, classical methods remain widely adopted for multi-center studies that require interpretable, deterministic outputs. Yet at cohort scale, many pipelines still estimate and apply the target appearance sequentially, leaving GPU parallelism underused.

GPU-capable packages such as Slideflow and torch-staintools already offer GPU-accelerated classical methods with a fit/transform API, while torchstain provides GPU-accelerated single-image normalization. High-throughput CPU systems (Anghel et al. 2019) and broader analysis platforms such as TIAToolbox (Pocock et al. 2022) likewise include classical stain normalization among wider workflows. What remains missing is a tool that combines a fit-once/transform-many API with optional fused CUDA kernels that coalesce per-method pixel operations into fewer GPU launches at batch scale. As WSI cohorts grow toward millions of slides (Zimmermann et al. 2024), normalization speed becomes the bottleneck.

Here we present StainX, a batch-oriented stain normalization framework with a two-stage fit/transform interface and an optional fused CUDA backend for NVIDIA GPUs. For Reinhard and histogram matching, source statistics are pooled over the batch, reducing per-tile overhead and ensuring a consistent appearance across tiles that share a batch.

## Implementation

StainX provides Histogram Matching, Macenko, and Reinhard normalization through a common fit/trans - form interface for batched tensors, with two execution backends: stainx[torch] and stainx[torch_cuda]. Macenko additionally exposes a fast CUDA mode (stainx[torch_cuda_fast]).

### Interface design

The fit method computes and stores method-specific parameters from the reference image(s) (Fig. 1a), namely three 256-bin channel histograms for Histogram Matching, three *L*^*^ *a*^*^ *b*^*^ (LAB) means and standard deviations for Reinhard, and a 3 × 2 stain matrix with two maximum concentrations for Macenko. Users should select a reference representative of the desired target appearance but StainX also includes a default reference derived from a representative H&E WSI. Users with different stain protocols or scan - ners may obtain better results with a custom reference. Fitted parameters can be saved and reused across runs. Subsequent transform calls reuse these parameters for the input batch, represented as (*N, C, H, W*) tensors for batch size, channels, height, and width, and return an output batch. Histogram Matching additionally accepts a channel-last (*N, H, W, C*) layout as well as the channels-first (*N, C, H, W*) layout. Reference parameters are computed through the PyTorch backend, while transformation uses the selected torch or torch_cuda backend. The provided StainNormalizerTransform module applies the same interface within PyTorch data pipelines.

**Figure 1:**
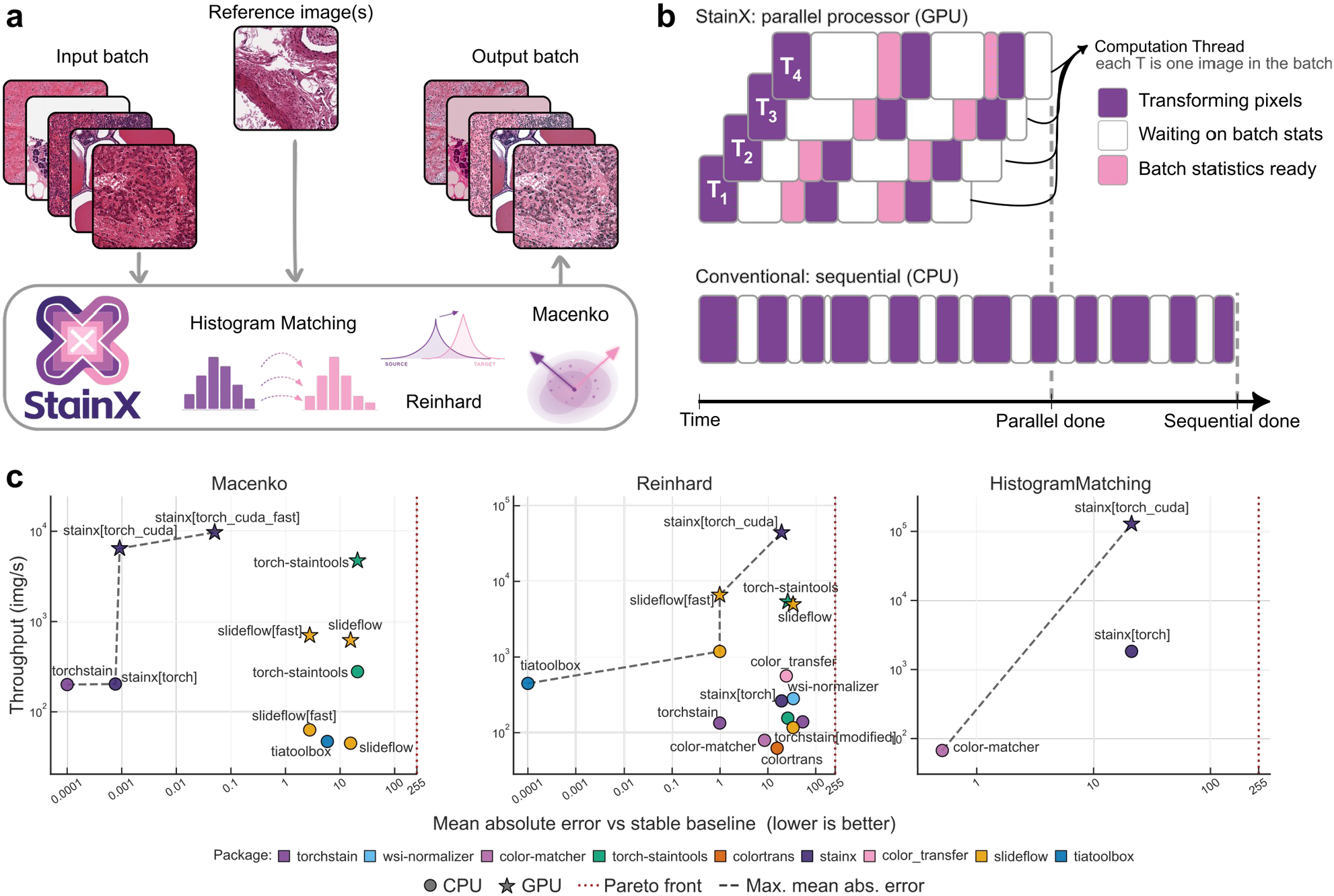
StainX design and benchmark evaluation. a) Given an input batch and reference image(s), StainX applies Histogram Matching, Reinhard, or Macenko to produce an output batch. b) StainX parallel processor runs one computation thread T per image in the batch (transforming pixels, waiting on batch stats, batch statistics ready), finishing at parallel done. Conventional CPU processes T _1_ … T _4_ sequentially until sequential done. c) Panels Macenko, Reinhard, and Histogram Matching show throughput (images/second) versus mean absolute error vs stable baseline (lower is better).

### Normalization methods

#### Histogram Matching

(Gonzalez and Woods 2009) aligns each source-channel intensity distribution with its reference distribution. Fit converts each reference channel into a normalized 256-bin histogram. Transform builds one source histogram and cumulative distribution function (CDF) per channel over all pixels in the input batch (not per image), locates reference quantiles by binary search, interpolates a 256-entry lookup table (LUT), and remaps the pixels. Consequently, each tile’s remapping depends on the composition of its batch.

#### Macenko

(Macenko et al. 2009) separates hematoxylin and eosin components in optical-density (OD) space. Unsigned 8-bit inputs are automatically converted to the [0, 1] range. StainX matches the torchstain convention (Barbano and Pedersen 2022), computing *O D* = − log ((255 *I* + 1) / *I*_0_) with *I*_0_ = 240, and retaining pixels for which *min* (*OD*) ≥ *β* = 0.15. The two leading eigenvectors of the resulting 3 × 3 covariance matrix define a stain plane, and its 1st and 99th angular percentiles (*α* = 1) determine the hematoxylin and eosin vectors. Fit stores these vectors as *H* ∈ ℝ^3 × 2^ together with the 99th-percentile stain concentrations. During transform, each image is decomposed in its own stain space because the OD mask contains a variable number of pixels. Its concentrations are scaled to the stored reference maxima and reconstructed with the reference *H*.

#### Reinhard

(Reinhard et al. 2001) matches channel statistics in LAB, transferring appearance from source to target in Fig. 1a. StainX converts red–green–blue (sRGB) through linear RGB and CIE XYZ under D65, then encodes floating-point OpenCV-compatible LAB (*L* × 2.55, with *a* and *b* offset by 128) rather than 8-bit quantized LAB. Fit stores channel means *μ*^*ref*^ and standard deviations *σ*^*ref*^ over reference pixels in that float encoding. Transform estimates source means 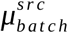 and standard deviations 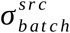 over all pixels in the input batch and applies 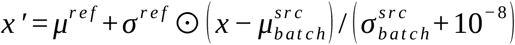 before conversion to sRGB. As with histogram matching, a tile’s output therefore depends on its batch-mates.

### GPU backend design

The portable torch backend is the primary contribution. It implements all methods with standard PyTorch tensor operations on any device PyTorch exposes without compiling an extension, including CPU, NVIDIA and AMD GPUs, and Apple Metal Performance Shaders (MPS). torch_cuda replaces the trans - form step of each normalizer with NVIDIA-specific fused CUDA kernels, and fit always uses the torch path. Fig. 1b contrasts this StainX parallel processor with a conventional CPU. Each computation thread T is one image in the batch and progresses through transforming pixels, waiting on batch stats, and batch statistics ready, reaching parallel done while the CPU path reaches sequential done later.

#### Histogram Matching

(CUDA transform). Each float tile is converted to 8-bit grey values. Many GPU blocks count how often each of the 256 grey levels appears in their chunk of the image in parallel, then those partial tallies are merged into one full histogram per channel. A CDF is built from the reference histogram, a 256-entry LUT maps each source grey level to its reference counterpart, and every pixel is remapped through that table in a single parallel pass before values are scaled and clamped back to the output range.

### Reinhard

(CUDA transform). All pixels are converted from sRGB to LAB in parallel. Source means and standard deviations over the batch are computed with PyTorch reductions. A single fused kernel sub - tracts the source statistics, scales by the reference standard deviation, and adds the reference mean for every pixel, after which a last kernel converts the normalized LAB back to sRGB.

### Macenko

(CUDA transform). One thread block is launched per image. Each block builds a tissue mask (pixels whose minimum OD exceeds *β*), accumulates a 3 × 3 OD covariance with warp-level reduction, and runs a closed-form analytic symmetric 3 × 3 eigensolver on thread 0 to obtain the hematoxylin and eosin stain-plane directions, all without leaving the GPU. Two precision modes are available: stable mode (stainx[torch_cuda]) uses float64 for the covariance and eigensolve with float32 downstream computations, and is recommended when numerical fidelity is critical; fast mode (stainx[torch_cuda_fast]) uses float32 throughout with float16 for large pixel tensors, and is intended for pipelines that maximize throughput. Both modes pass the correctness suite against the reference. Because Macenko’s per-image eigendecomposition is not trivially batch-parallel, its CUDA speedup over stainx[torch] is substantial but more modest than for Reinhard or Histogram Matching.

Users may manually select a backend, but automatic selection is implemented by using torch_cuda when an NVIDIA CUDA device and its compiled extension are available.

## Benchmarking

### Setup

We benchmarked StainX against established tools to assess real-world throughput and numerical fidelity. Experiments used batches of 128 tiles with size of 256 × 256 pixel RGB, with all timing measured directly on the device to exclude host–device transfers. Each transformation was timed over 100 runs following 30 warm-up iterations. Benchmarks were run on a Lambda Vector workstation with an AMD Threadripper Pro 5975WX (32 cores) and an NVIDIA RTX A6000 (48 GB). Mean absolute error was computed against reference implementations: torchstain (Barbano and Pedersen 2022) and scikit-image (Van der Walt et al. 2014). The surveyed GPU-capable peers included slideflow (Dolezal et al. 2024), torch-staintools (Zhou 2024), torchstain, tiatoolbox (Pocock et al. 2022), and color-matcher (Hahne 2020), while the CPU packages are wsi-normalizer (Cui 2025), colortrans (Steinberg 2026), and color_transfer (Rosebrock 2014).

### Results

StainX led across all three methods (Fig. 1c). Relative to its own portable torch CPU backend, the fused CUDA path achieved 168 × the throughput for Reinhard, 70 × for Histogram Matching, and 32 × for Macenko (rising to 48 × in the fast precision mode). Against the fastest GPU peers, Reinhard throughput was about 7–8 × that of Slideflow and torch-staintools, Macenko’s fast mode doubled torch-staintools throughput at comparable accuracy, and Histogram Matching had no comparable GPU peer. For Reinhard and histogram matching, peer packages estimated source statistics per image whereas StainX pooled them over the batch, so peer speedups were not like-for-like. Numerical correctness was verified against implementations: outputs matched within at most one grey level for Reinhard and Histogram Matching compared to torchstain and scikit-image, and within two grey levels for Macenko against torch-stain (mean absolute error ≤ 0.35). These results demonstrate that StainX delivers reference-quality normalization at the throughput required for large-scale preprocessing.

## Conclusion

StainX makes reference-controlled stain normalization practical at cohort scale, delivering up to 168 × CPU speedup and 7–8 × the throughput of the fastest GPU peers for Reinhard, while matching reference implementations within 1–2 grey levels. The library is pip-installable across Linux, macOS, and Windows, with a documented API, continuous integration testing, and a portable PyTorch backend that requires no CUDA compilation. Normalization quality depends on reference image selection; we recommend visual spot-checking of a subset of normalized tiles in any cohort-scale run.

## Acknowledgements

We thank the IT team at CeMM for access and maintenance of the CeMM HPC cluster.

## Author contributions

S.M. designed the project, implemented StainX and wrote the manuscript draft, Y.Z. contributed to the validation of the implementations, and A.R. supervised the research. All authors revised the manuscript.

## Supplementary material

No supplementary material is provided for this article.

## Data availability

The current release of StainX (v0.1.3) is available at https://github.com/rendeirolab/stainx/ under the GNU GPL v3.0 or later and installable via `pip install stainx` from PyPI. The portable PyTorch backend requires no CUDA compilation and works on any device PyTorch supports (Linux, macOS and Windows, Python ≥3.11, PyTorch ≥2.0), while the optional CUDA backend is compiled on first use for NVIDIA GPUs. Tagged releases will be archived on Zenodo (DOI to be added upon deposition). Bench - mark scripts, logs, and example tiles used for Fig. 1c are included in the repository.

## References

Anghel, Andreea, Milos Stanisavljevic, Sonali Andani, Nikolaos Papandreou, Jan Hendrick Rüschoff, Peter Wild, Maria Gabrani, and Haralampos Pozidis. 2019. “A High-Performance System for Robust Stain Normalization of Whole-Slide Images in Histopathology.” Frontiers in Medicine 6: 193. 10.3389/fmed.2019.00193.

Barbano, Carlo Alberto, and André Pedersen. 2022. “Torchstain (Version 1.4.1).” Zenodo. 10.5281/zenodo.6979540.

Chen, Richard J, Tong Ding, Ming Y Lu, Drew FK Williamson, Guillaume Jaume, Andrew H Song, Bowen Chen, et al. 2024. “Towards a General-Purpose Foundation Model for Computational Pathology.” Nature Medicine 30 (4): 1057–73. 10.1038/s41591-024-02857-3.

Cui, Haoyu. 2025. “Wsi-Normalizer (Version 1.0.1).” PyPI. https://pypi.org/project/wsi-normalizer/1.0.1/.

Dolezal, James M., Sara Kochanny, Emma Dyer, Siddhi Ramesh, Andrew Srisuwananukorn, Matteo Sacco, Frederick M. Howard, and et al. 2024. “Slideflow: Deep Learning for Digital Histopathology with Real-Time Whole-Slide Visualization.” BMC Bioinformatics 25 (1): 134. 10.1186/s12859-024-05758-x.

Gonzalez, Rafael C, and Richard E Woods. 2009. Digital Image Processing. 3rd ed. Pearson Education. https://search.worldcat.org/search?q=bn:9780131687288.

Hahne, Christopher. 2020. “Color-Matcher (Version 0.6.0).” PyPI. https://pypi.org/project/color-matcher/0.6.0/.

Macenko, Marc, Marc Niethammer, James S Marron, David Borland, John T Woosley, Xiaojun Guan, Charles Schmitt, and Nancy E Thomas. 2009. “A Method for Normalizing Histology Slides for Quantitative Analysis.” In 2009 IEEE International Symposium on Biomedical Imaging: From Nano to Macro, 1107–10. IEEE. 10.1109/ISBI.2009.5193250.

Pocock, Johnathan, Simon Graham, Quoc Dang Vu, Mostafa Jahanifar, Srijay Deshpande, Giorgos Hadjigeorghiou, Adam Shephard, et al. 2022. “TIAToolbox as an End-to-End Library for Advanced Tissue Image Analytics.” Communications Medicine 2 (1): 120. 10.1038/s43856-022-00186-5.

Reinhard, Erik, Michael Adhikhmin, Bruce Gooch, and Peter Shirley. 2001. “Color Transfer Between Images.” IEEE Computer Graphics and Applications 21 (5): 34–41. 10.1109/38.946629.

Rosebrock, Adrian. 2014. “Color_transfer (Version 0.1).” PyPI. https://pypi.org/project/color_transfer/0.1/.

Steinberg, Daniel. 2026. “Colortrans (Version 1.1.0).” PyPI. https://pypi.org/project/colortrans/1.1.0/.

Tellez, David, Geert Litjens, Péter Bándi, Wouter Bulten, John-Melle Bokhorst, Francesco Ciompi, and Jeroen Van Der Laak. 2019. “Quantifying the Effects of Data Augmentation and Stain Color Normalization in Convolutional Neural Networks for Computational Pathology.” Medical Image Analysis 58: 101563. 10.1016/j.media.2019.101563.

Van der Walt, Stefan, Johannes L Schönberger, Juan Nunez-Iglesias, François Boulogne, Joshua D Warner, Neil Yager, Emmanuelle Gouillart, and Tony Yu. 2014. “Scikit-Image: Image Processing in Python.” PeerJ 2: e453. 10.7717/peerj.453.

Zhou, Yufei. 2024. “Torch-Staintools (Version 1.0.7).” Zenodo. 10.5281/zenodo.10453806.

Zimmermann, Eric, Eugene Vorontsov, Julian Viret, Adam Casson, Michal Zelechowski, George Shaikovski, Neil Tenenholtz, et al. 2024. “Virchow2: Scaling Self-Supervised Mixed Magnification Models in Pathology.” arXiv Preprint arXiv:2408.00738. 10.48550/arXiv.2408.00738.

